# Graphene Oxide as 2D Platform for Complexation and Intracellular Delivery of siRNA

**DOI:** 10.1101/486522

**Authors:** Irene de Lázaro, Sandra Vranic, Domenico Marson, Artur Filipe Rodrigues, Maurizio Buggio, Adrián Esteban-Arranz, Mariarosa Mazza, Paola Posocco, Kostas Kostarelos

## Abstract

The development of efficient and safe nucleic acid delivery vectors remains an unmet need holding back translation of gene therapy approaches into bedside. Graphene oxide (GO) could help bypass such bottleneck thanks to its large surface area, versatile chemistry and biocompatibility, which could overall enhance transfection efficiency while abolishing some of the limitations linked to the use of viral vectors. Here, we aimed to assess the capacity of bare GO, without any further surface modification, to complex a short double-stranded nucleic acid of biological relevance (siRNA) and mediate its intracellular delivery. GO formed stable complexes with siRNA at 10:1, 20:1 and 50:1 GO:siRNA mass ratios. Complexation was further corroborated by atomistic molecular dynamics simulations. GO:siRNA complexes were promptly internalized in a primary mouse cell culture, as early as 4 h after exposure. At this time point, intracellular siRNA levels were comparable to those provided by a lipid-based transfection reagent that achieved significant gene silencing. Time-lapse tracking of internalized GO and siRNA evidenced a sharp decrease of intracellular siRNA from 4 to 12 h, while GO was sequestered in large vesicles, which may explain the lack of biological effect (i.e. gene silencing) achieved by GO:siRNA complexes. This study underlines the potential of non-surface modified GO flakes to act as 2D siRNA delivery platforms, without the need for cationic functionalization, but warrants further vector optimization to allow effective release of the nucleic acid and achieve efficient gene silencing.

---

Engineering efficient, targeted and safe nucleic acid delivery vectors that are also easily scalable and reproducibly manufactured remains a challenge to be tackled if promising gene therapy strategies, including gene silencing and editing, among others, are to become a clinical reality. Viral vectors outperform non-viral counterparts in gene transfer efficiency and consequently have been both much more popular and successful in clinical trials to date^1^, but are yet to completely overcome packaging restrictions and large-scale production constraints, in addition to their controversial safety profile^2^. On the other hand, promising developments by non-viral carriers — including lipid, polymeric nanoparticles, and carbon nanotubes, among others — to circumvent some of such limitations have been overshadowed by low transfection efficiencies and poor *in vitro*-to-*in vivo* translation^3^.

2-dimensional (2D) nanomaterials are attracting increasing attention in many areas of research, including biomedicine. As molecular transporters, they could make a difference in cargo loading and transfer efficiencies thanks to their high surface-to-volume aspect ratio, much larger than that of other nanomaterials^4^. Graphene oxide (GO) offers further advantages based on the presence of multiple oxygen functionalities, including high aqueous dispersibility, thus compatibility with the biological milieu, and facile chemical functionalization. The latter can be exploited to favor interactions with molecules of diverse chemical nature^5^. GO sheets are also known to internalize within phagocytic and non-phagocytic cells, although uptake mechanisms are not yet fully understood and are clearly impacted by the specific lateral dimensions and number of layers of the material^6-7^. Finally, its low cytotoxicity, compared to that of other nanomaterials, further encourages the biomedical applications of GO. Indeed, no adverse effects have been reported upon *in vivo* administration of this material following various routes, which included intravenous, intraperitoneal and pulmonary administration^8^.

Despite these advantages, the performance of GO in the specific context of nucleic acid delivery could be hindered by unfavorable electrostatic interactions led by the presence of negative charges in both vector and cargo. This is especially relevant when the oligonucleotide of choice is double stranded, since hydrophobic and π-π interactions between the nucleobases and the GO lattice are sterically hindered^6^. A number of studies (reviewed elsewhere^9^) have used GO to complex and deliver double-stranded nucleic acids intracellularly, including plasmid DNA (pDNA) and small interfering RNA (siRNA). However, all of them relied on functionalization of the material with positively charged moieties ― including cationic polymers^10^, polysaccharides^11^ and cell penetrating peptides^12^ ― which have previously been used as delivery vectors on their own, but whose biocompatibility is far from ideal. GO was therefore used as one of the components of more complex delivery systems in such studies. Whether bare GO flakes, without further functionalization, can complex and deliver double-stranded oligonucleotides still remains unknown.

In this study, we aimed to investigate whether bare GO could interact with a short, double-stranded nucleic acid of biological relevance, siRNA, and act as its efficient intracellular delivery vector without further functionalization. To address this, we used endotoxin-free, small GO flakes (mean lateral size < 1µm) obtained synthetically by a modified Hummer’s method^13-14^ and assessed complexation at different GO:siRNA mass ratios, both experimentally and via molecular dynamics (MD) simulation. We interrogated the intracellular fate of both components, carrier and cargo, in a primary mouse cell culture system, as well as the capacity of the oligonucleotide to desorb from the GO lattice and induce gene silencing. The performance of bare GO as siRNA delivery vector was compared to that of a benchmark lipid-based transfection reagent (Lipofectamine^®^).

## Results

### Bare GO complexes siRNA

Endotoxin-free GO sheets were synthesized *in house* following a modified Hummers method^13-14^ (see Experimental). A summary of the physicochemical characterization of the GO dispersion in water, reported elsewhere^15^, is provided in **Table S1**. In brief, structural properties were studied by transmission electron microscopy (TEM) and atomic force microscopy (AFM), showing that the lateral dimensions of GO flakes ranged between 0.05 - 2 µm. Flake thickness corresponded to 1-2 GO layers, approximately. A representative AFM image is provided in **Figure S1**.

We first investigated the capacity such flakes, without any further functionalization with cationic moieties, to interact with a short (23 bp long) double-stranded nucleic acid (siRNA) by directly mixing the carbonaceous material (dispersed in water) with the nucleic acid and assessing the electrophoretic mobility of the latter. A progressive decrease in the intensity of the band corresponding to free (non-complexed) siRNA, which correlated positively with the increase in GO:siRNA mass ratio, indicated that the material was able to interact with siRNA at all mass ratios under investigation —10:1, 20:1 and 50:1 — and that complexation improved with the increased amount of GO (Figure 1a). GO quenches fluorophores that interact with its sp^2^ aromatic domain via fluorescence resonance energy transfer (FRET) and dipole–dipole coupling effects, which explains its popularity in the biosensing field^16-17^. Gelred™ is a commercially available oligonucleotide dye that emits fluorescence at 620 nm upon interaction with single-stranded DNA (ssDNA), double-stranded DNA (dsDNA) or RNA. Direct recording of the fluorescence spectrum emitted by this molecule after incubation with siRNA or GO:siRNA complexes showed a decrease in fluorescence intensity in the presence of GO. Again, we found a positive relationship between the increasing GO:siRNA mass ratio and the efficiency of complexation (i.e. decrease in fluorescence intensity) (Figure 1b).

**Figure 1.**
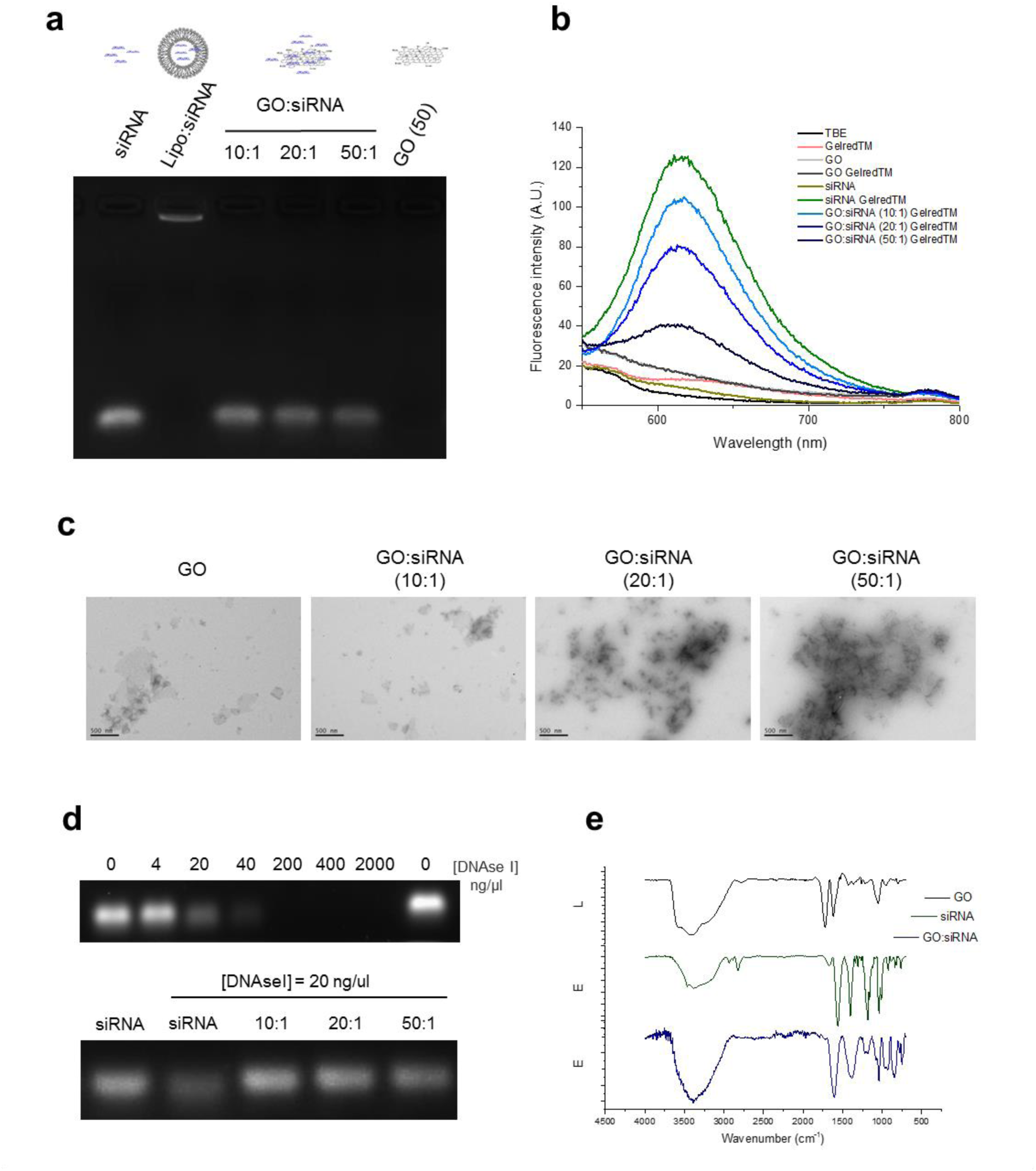
GO:siRNA complexation studies. Complexation of double-stranded siRNA was investigated at 10:1, 20:1 and 50:1 GO:siRNA mass ratios by **(a)** gel retardation assay, **(b)** quenching of the fluorescence signal of a nucleic acid dye (Gelred™) and **(c)** transmission electron microscopy (TEM). **(d)** Exposure to DNAseI confirmed complexation, given the absence of siRNA degradation in the presence of GO. (The top gel shows degradation of free siRNA by increasing concentrations of DNAseI, the bottom gel shows degradation of free or complexed siRNA at 20 ng/µl DNAseI). **(e)** Differences in ATR-FTIR spectra of GO, siRNA and GO:siRNA (50:1 mass ratio) confirmed the interaction of the oligonucleotide with the GO surface.

In agreement, a denser siRNA cloud was observed in transmission electron microscopy (TEM) micrographs of the complexes with the increase in GO:siRNA mass ratio (Figure 1c). Of note, after complexation, samples were diluted to reach equal GO concentrations to avoid bias in the analysis of the images. GO:siRNA exposure to DNAseI, at a concentration known to induce siRNA degradation (Figure 1d, top gel), provided further evidence of GO:siRNA interaction. When complexed to GO, siRNA was not degraded by the enzyme (Figure 1d, bottom gel). The same effect has been described by others using GO:ssDNA complexes, and may be explained by the GO lattice sterically impeding access of the enzyme to the nucleic acid^16, 18^. Finally, the ATR-FTIR spectrum of GO:siRNA (50:1 mass ratio) significantly differed from that of GO alone, and further confirmed the contribution of the oligonucleotide on the GO surface (Figure 1e). First, the stretching vibration band corresponding to -C=C unsaturated bonds in GO shifted from 1619 to 1605 cm^-1^, which may be due to interactions with the α-helical structure of the nucleic acid. Most strikingly, the band referred to stretching vibrations of –C=O bonds in carboxylic groups (1721 cm^-1^) disappeared upon complexation, which suggests that such groups could contribute to the formation of hydrogen bonds driving siRNA adsorption.

siRNA complexation was also supported by molecular-level theoretical predictions. MD studies of adsorption of a siRNA double strand on a GO flake were carried out in aqueous environment and at different ionic strengths, namely 0, 150, and 200 mM NaCl. We found that the oligonucleotide could be fully and stably adsorbed near the GO interface in 2 out of 6 and 6 out of 6 independent runs considered at 0 mM and 150/200 mM NaCl, respectively **(Video S1 and Video S2)**. Representative snapshots of the equilibrated GO:siRNA complexes are shown in Figure 2a-b and **Figure S2**. To gain insights into the GO:siRNA binding affinity, steered MD (SMD) experiments of siRNA pulled away from the GO surface were then performed starting from equilibrated complexes. The force required to unbind the oligonucleotide was monitored as a function of the distance from the center of mass of GO and siRNA (Figure 2c). The profiles obtained indicated the substantial influence of the salt concentration on the binding affinity, with rupture forces (i.e. the maximum force required to unbind siRNA from GO) decreasing as NaCl content decreased. At 200 nM NaCl, the peak corresponded to circa 4060 pN, but decreased to 2520 pN and 840 pN at 150 and 0 mM NaCl, respectively. This observation may be explained by the alleviation of coulombic repulsions, triggered by negative charges from oxygen functionalities on the GO surface and from phosphate groups in the siRNA, in the presence of complementary ions. Indeed, the addition of salts to the solution brought the double strand close to the GO surface (Figure 2a-b and Figure S2), where favorable short-range polar interactions can occur. This hypothesis is also in line with the evidence obtained by ATR-FTIR as described above.

**Figure 2.**
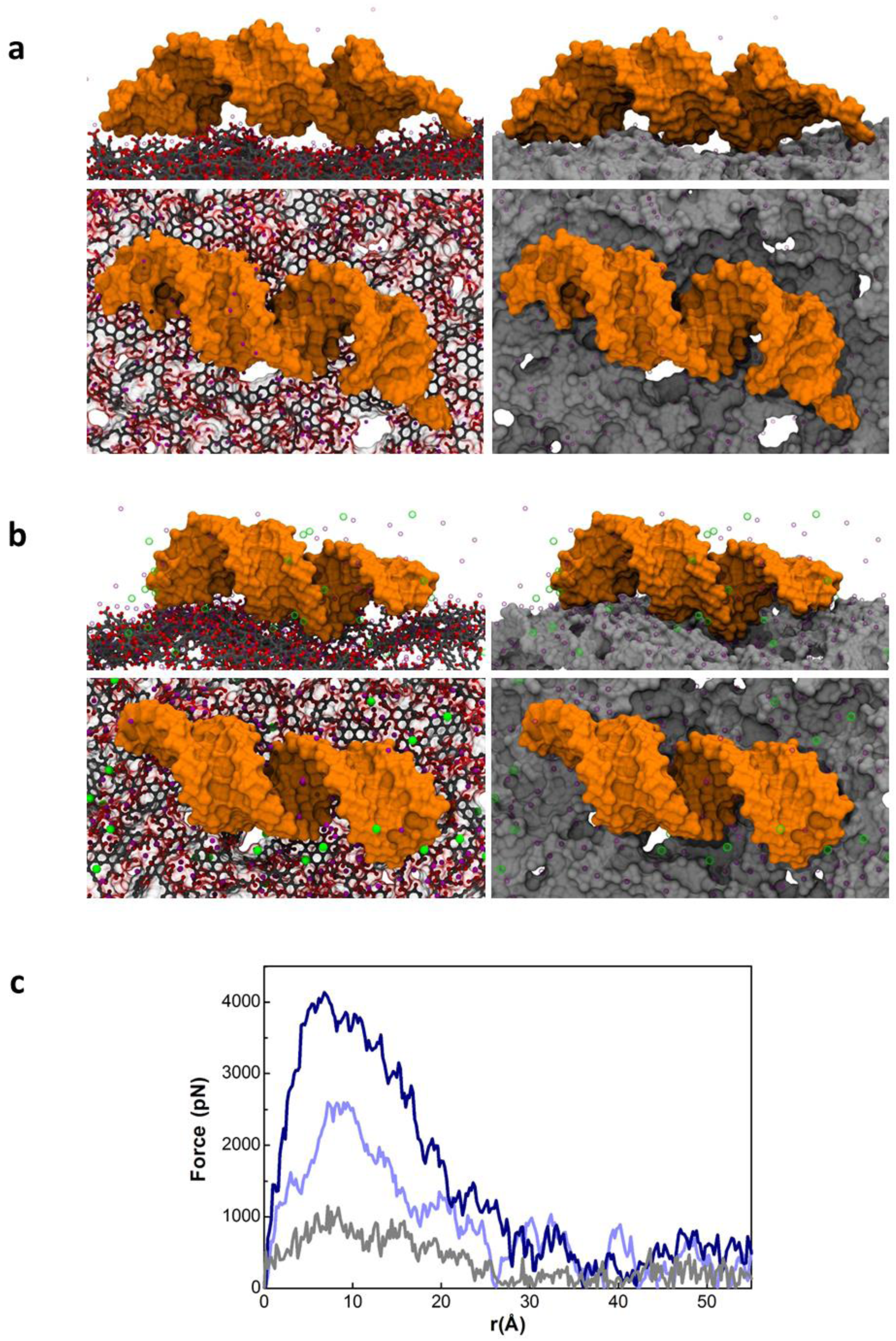
GO:siRNA binding/unbinding by MD simulation. Snapshots extracted from MD calculation of siRNA (represented as an orange shaded surface) in complex with a GO sheet **(a)** without complementary salts and **(b)** at 150 mM NaCl ionic strength (top panels, lateral view; bottom panels, top view). Images on the left show the chemical structure of GO explicitly represented with carbon and oxygen atoms colored in grey and red, respectively. In the right panels, we depicted the GO sheet as a grey surface to highlight 3D features. For clarity, water is not shown and Na^+^ and Cl^-^ ions are visualized as purple and green spheres, respectively. **(c)** SMD simulation of the unbinding process of siRNA from GO. The pulling force as a function of the distance from the center of mass of GO and siRNA in the direction perpendicular to the GO plane is monitored. Color legend: 0 mM, grey; 150 mM, light blue; 200 mM, dark blue.

### GO:siRNA complexes internalize in primary mouse cells without inducing cytotoxicity

Driven by our encouraging results on GO:siRNA complexation, and by other studies that have reported efficient cellular uptake of GO even in non-phagocytic cells (reviewed elsewhere^7^), we aimed to investigate whether GO could be used as an intracellular siRNA delivery vector. We first examined any cytotoxic responses to GO:siRNA complexes on a primary mouse cell culture system. A GO material similar to the one utilized here — differing only in the graphite source, but with practically identical physicochemical properties — did not induce cytotoxicity in a human lung epithelial cell line (BEAS-2B) at the doses included in this study^19^. Here, mouse embryonic fibroblasts (MEFs) obtained from CD1 mice were exposed to GO:siRNA complexes, where siRNA was a scrambled sequence known to be inert to cells. Even with the highest mass ratio tested (50:1) cell morphology remained unaltered 24 h after treatment **(Figure S3a)** and no changes were detected in the percentage of live cells assessed by AnnexinV/PI staining using flow cytometry **(Figure S3b)**.

Next, we followed the internalization of GO and siRNA in MEFs. Thanks to the intrinsic fluorescence of the material, GO uptake can be monitored by confocal microscopy without further functionalization or labelling, as demonstrated in a previous study from our laboratory^19^. However, fluorophore quenching impeded the visualization of a fluorescently-labelled siRNA complexed onto GO (data not shown). Instead, we set out to address the uptake of carrier and cargo separately and to design a PCR-based strategy to quantify internalization of the latter. Live confocal microscopy evidenced cellular internalization of the GO component in GO:siRNA complexes, as early as 4 h after exposure (Figure 3a and 3b), where siRNA was a non-coding, non-fluorescent sequence. GO uptake was dose and time dependent. Of note, in the treatment with different GO:siRNA mass ratios, siRNA concentration was kept constant while GO changed accordingly. GO intracellular uptake was significantly higher 24 h after treatment, but was only observed when MEFs were exposed to GO:siRNA complexes in serum-free conditions for the first 4 h **(Figure S4)**. Impaired cellular internalization in the presence of serum proteins has been reported for other delivery vectors^20-21^, including benchmark transfection reagents such as Lipofectamine^®22^, and was evidenced here by the absence of siRNA intracellular delivery (Figure 3c and Figure S5a) and gene knockdown **(Figure S5b)** when the lipid-based vector was used in the presence of FBS.

**Figure 3.**
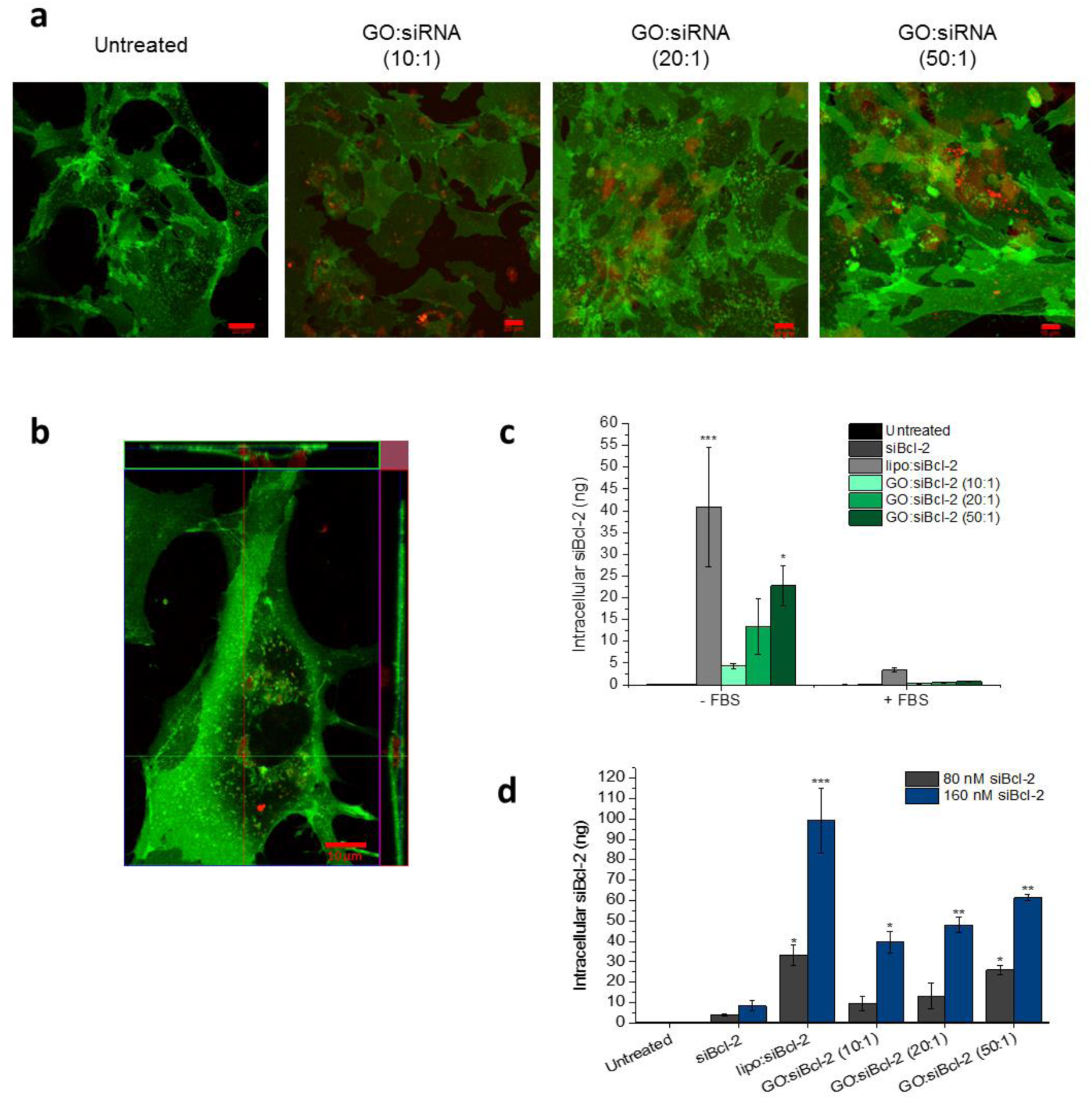
Uptake of GO:siRNA complexes 4 h after exposure. **(a)** GO uptake in MEFs was visualised by live cell confocal microscopy, 4 h after transfection, thanks to the intrinsic fluorescence of the material. Images represent maximum intensity projection of the fluorescence across the Z-stack, corresponding to plasma membrane staining (green) and GO (red). Scale bars show 10 or 20 µm. **(b)** Orthogonal view confirms intracellular localisation of the GO, upon exposure to GO:siRNA (50:1). Green signal corresponds to cell membrane staining, red signal corresponds to GO. Scale bar shows 10 µm. **(b)** Intracellularly delivered siRNA was quantified by stem-loop PCR in the presence or absence of FBS (80 nM siRNA) and **(c)** comparing two different siRNA concentrations (80 vs 160 nM) in the absence of FBS.*p<0.05, **p<0.01 and ***p<0.001 represent statistically significant differences in siBcl-2 levels compared to untreated cells. No statistically significant differences were found between lipo:siRNA [80 nM] and GO:siRNA [160 nM] groups (one-way ANOVA, Tukey post-hoc test, n=3).

To confirm siRNA internalization, we focused on the 4 h time point, as the earliest time point when GO was detected intracellularly. GO:siRNA complexes were formed with a siRNA sequence specific to the *Bcl-2* gene (si-Bcl2) and specific primers were designed to amplify the oligonucleotide via stem-loop PCR. We took advantage of the complete blockade of lipo:siRNA and GO:siRNA uptake in the presence of FBS to confirm that this method detected intracellularly delivered siBcl-2 exclusively, and not the oligonucleotides that could remain in complexes attached to the cell membrane. Indeed, negligible siBcl-2 levels were detected when exposure took place in serum-containing medium (Figure 3c). On the contrary, and in agreement with our complexation studies, increasing amounts of siBcl-2 were detected intracellularly with the increase in GO:siBcl-2 mass ratio, when complexes were incubated in serum-free medium (Figure 3c). However, none of the mass ratios tested outperformed Lipofectamine^®^. We therefore decided to investigate if higher levels of intracellular siBcl-2 could be attained by increasing the initial concentration of the nucleic acid. We compared two different siBcl-2 concentrations, 80 and 160 nM, but fixed the mass ratios under investigation. Starting at [siBcl-2]= 160 nM, all GO:siBcl-2 mass ratios tested delivered as much intracellular siBcl-2 as Lipofectamine^®^ did with an initial [siBcl-2]= 80 nM (Figure 3d).

### GO:siRNA complexes fail to release functionally active siRNA

To investigate if siBcl-2 was released from the GO lattice in the cell cytosol, while retaining functional integrity, we first performed gene knockdown studies in MEFs. Both siBcl-2 concentrations investigated in quantification studies, 80 and 160 nM, were tested. Exposure to GO is known to induce changes in gene expression^23^, even in genes whose expression is typically stable and that are commonly used as references for RT-qPCR normalization^24^. Therefore, we tested the stability of ten accepted “reference” genes under the specific experimental conditions of our study and concluded that use of the two most stable (*Tbp* and *Hmbs*) allowed reliable normalization of gene expression data **(Figure S6)**. The siBcl-2 sequence utilized in this study was functionally verified via transfection with lipo:siBcl-2 complexes, which induced significant downregulation of the target mRNA 24 h after treatment, at both 80nM and 160 nM siBcl-2 starting concentrations (63% and 76% knockdown, respectively). Surprisingly, none of the GO:siBcl-2 conditions tested induced *Bcl-2* downregulation, even at the highest siBcl-2 concentration (Figure 4a and 4b). This observation suggested that siRNA might not be efficiently released from the GO lattice, or functionally active, upon cellular internalization.

**Figure 4.**
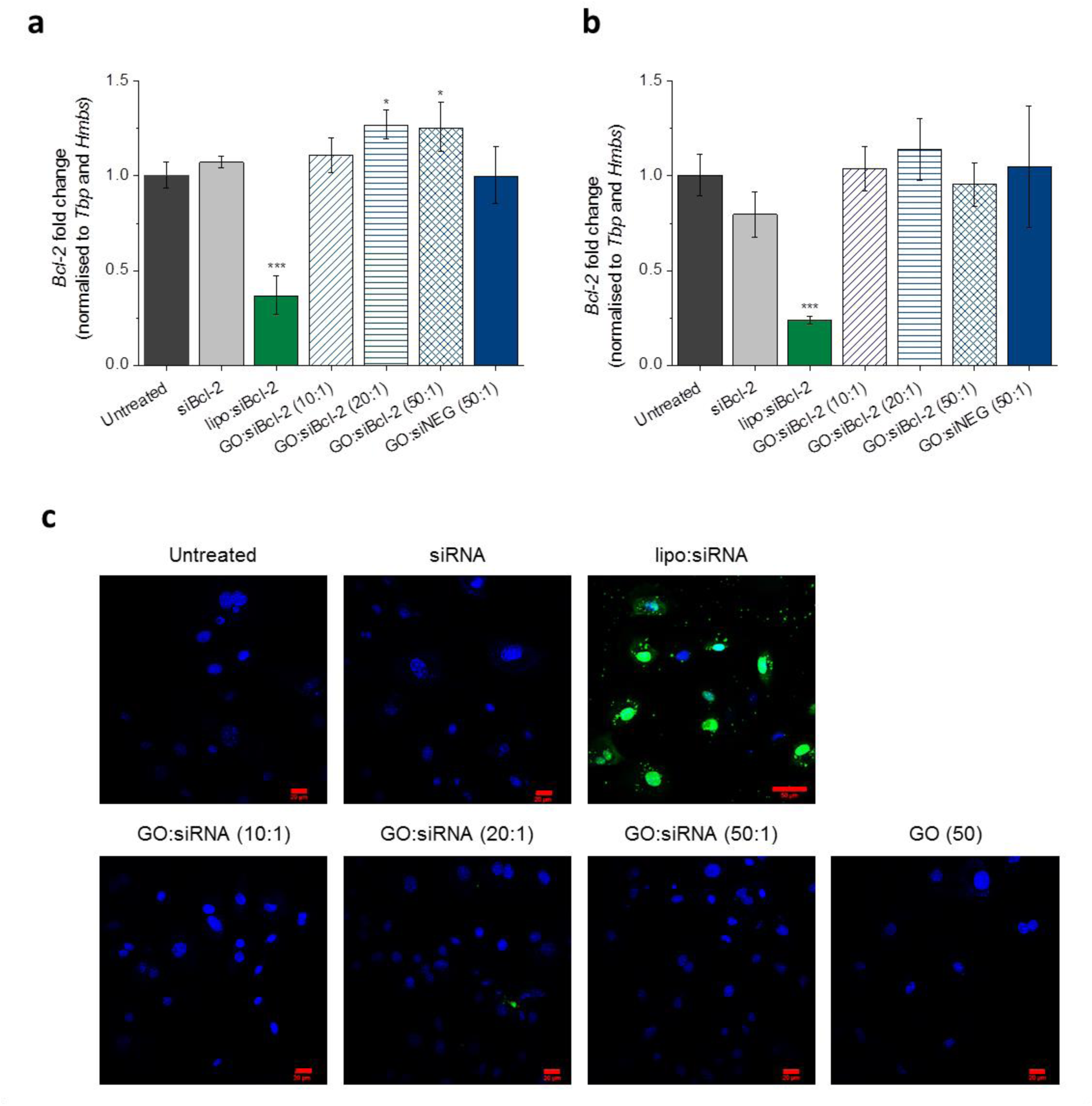
In vitro functional assessment of GO:siRNA complexes. *Bcl-2* mRNA expression was studied by RT-qPCR 24 h after transfection with GO:siBcl-2 complexes at **(a)** [siBcl-2]= 80 nM and **(b)** [siBcl-2]=160 nM. *p<0.05 and ***p<0.001, one-way ANOVA and Tukey post-hoc test, n=3. **(c)** siRNA desorption from GO and intracellular trafficking were studied with a RISC-independent, nuclear-targeted and fluorescently-labelled siRNA (siGLO) complexed to GO. 4 h after transfection, siGLO only localised to the nucleus when delivered by a lipid-based transfection reagent. No fluorescence was recovered in any of the GO:siGLO conditions. Green signal corresponds to 6-FAM labelling of siGLO oligonucleotide, blue signal corresponds to Hoechst nuclear staining. Scale bar shows 20 or 50 µm.

To investigate potential pitfalls in siRNA desorption, siGlO, a RISC-independent non-coding siRNA that contains a nuclear localization signal (NLS) sequence and is conjugated to 6-carboxyfluorescein (6-FAM), was used. GO:siGLO complexation was confirmed by 6-FAM quenching and, as observed with unmodified siRNA, improved with the increase in GO:siGLO mass ratio **(Figure S7a).** We hypothesized that if the oligonucleotide was able to detach from the carbon lattice intracellularly and traffic to its site of action, green fluorescence would be recovered in the cell nucleus. 4 h after transfection with lipo:siGLO or GO:siGLO complexes, bright green fluorescence co-localized with nuclear Hoechst staining only in cells transfected with Lipofectamine^®^ (Figure 4c). Even at a later time point, 24 h after transfection, siGLO fluorescence was still not recovered from GO:siGLO complexes **(Figure S7b)**, which suggested that the oligonucleotide was not able to dissociate from the GO surface and traffic intracellularly.

### Intracellular siRNA levels decrease rapidly after transfection while GO is sequestered in distinct intracellular vesicles

Moved by the lack of siRNA biological activity and trafficking inside the cell, we aimed to investigate the intracellular fate of GO and siRNA over time. Quantification of intracellular siBcl-2 at various time points after exposure indicated that the lipid-based transfection reagent Lipofectamine^®^ was able to sustain siBcl-2 levels for at least 24 h. In contrast, intracellular siBcl-2 sharply decreased from 4 to 12 h when delivered in GO:siBcl-2 complexes (Figure 5a). Such a striking difference in the intracellular persistence of the oligonucleotide is likely to be the reason behind the different performance of both vectors to silence the mRNA target. The same trend in siBcl-2 intracellular kinetics was found in a different cell type, the murine breast cancer cell line 4T1, which suggested that rapid siBcl-2 depletion upon GO:siBcl-2 internalization is not exclusive to MEFs **(Figure S8a)**.

**Figure 5.**
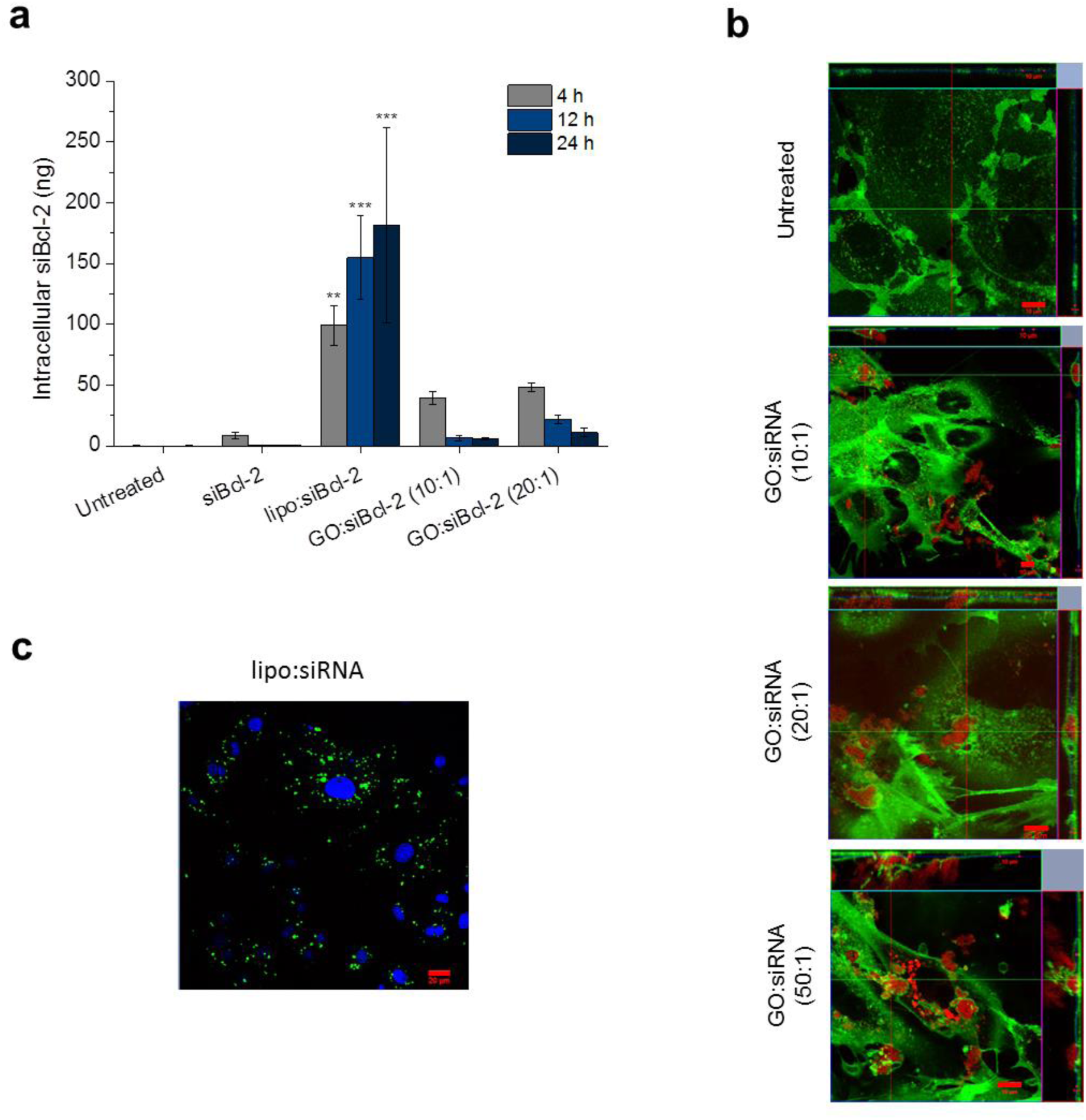
Intracellular fate of GO:siRNA complexes. **(a)** PCR-based quantification of intracellularly delivered siRNA 4, 12 and 24 h after transfection indicated a sharp decrease of intracellular siRNA between 4 and 12 h time points when delivered by GO:siRNA complexes (**p<0.01 and ***p<0.001, one-way ANOVA and Tukey post-hoc test, n=3). **(b)** 24 h after transfection, GO was still present intracellularly, but sequestered in large vesicles, as shown by live confocal microscopy. Green – Cell Membrane Mask, Red – GO. Scale bars show 10 or 20 µm. **(c)** Confocal microscopy image of MEFs treated with lipo:siRNA complexes, where siRNA was fluorescently labelled, 24h after transfection. Green – non-coding siRNA-488, Blue – nucleus (Hoechst). Scale bar shows 20 µm.

The intracellular fate of GO was monitored over the same timeframe via confocal microscopy. Starting from 4 h after the initial exposure, GO was found to accumulate in large intracellular vesicles that became more obvious with time (Figures 3b and S7b). The material remained sequestered in such structures that did not traffic inside or outside the cell for over 24 h (Figure 5b, Figure S8c and Video S3), when the amount of intracellular siBcl-2 detected was negligible (Figure 5a). Together, lack of evidence of siRNA desorption from the GO lattice, rapid decrease of its intracellular levels upon internalization and persistence of the material inside vesicles, suggest that the oligonucleotide could be trapped and degraded within such intracellular structures. In cells treated with lipo:siRNA complexes, siRNA was localized in small endocytic vesicles characteristic of liposomal-mediated delivery (Figure 5c), which were morphologically very different from those formed in the presence of GO (Figure 5b). Of note, this pattern of intracellular uptake was also found when cells were exposed to GO alone and therefore was not triggered by the coating with siRNA double strands **(Figure S8d)**. This observation suggested that the different performance of the liposome and GO vectors may not only be related to the release of the oligonucleotide from the carrier, but also to the mechanism governing the uptake of the complexes.

## Discussion

We show here that GO, without further functionalization, forms stable complexes with a short double-stranded nucleic acid (siRNA) (Figures 1-2, Figure S2 and Videos S1-S2). The majority of studies aiming to elucidate and exploit potential interactions between GO and nucleic acids focused on single-stranded oligonucleotides, given the higher availability of the nucleobases to interact with the GO lattice. Indeed, hydrophobic and π-π interactions between the bases and the aromatic domain of GO are known to drive the adsorption of single strands on the material^25-26^, counteracting electrostatic repulsion triggered by the presence of negative charges in both components. Accordingly, studies by Lu *et al* and He *et al* reported the binding affinity of ssDNA for GO to be significantly higher than that of dsDNA^27-28^. In this context, GO has been used to design intracellular biosensors in which single-stranded aptamers or molecular beacons specific to intracellular targets are adsorbed onto the GO lattice^16, 18^. However, other studies suggested that nucleobases at the double-strand terminus could be available to establish π-π stacking interactions ^29^, or that these could be formed upon partial deformation of the double helix on the GO surface^30^. In addition, contribution of hydrogen bonds between oxygen groups in GO and polar groups in the nucleotide strand has recently been accounted^31^. Our ATR-FTIR data supports this observation, given the obvious disappearance of the band corresponding to -C=O bonds from carboxyl groups in GO (Figure 1e).

These findings suggest opportunities to utilize GO as an intracellular nanocarrier of double-stranded oligonucleotides which can be used in applications other than biosensing, including gene silencing and forced expression of specific gene products. In fact, GO has already been used in a number of studies to load and deliver double-stranded plasmid DNA and siRNA, albeit always as a component of more complex delivery systems^9^. As such, GO had no direct interaction with the nucleic acid but provided a scaffold for cationic moieties — such as positively charged polymers^10^, dendrimers^32^, polysaccharides^11^ and peptides^12^— that complexed negatively charged oligonucleotides through electrostatic interactions. In fact, most of such materials can be used as gene delivery vectors on their own, and have been exploited as such for years, but often limited by cytotoxic effects elicited by the abundance of positive charges^33-34^. GO contributed in ameliorating such cytotoxic character, for example, by allowing the reduction of the molecular weight of the cationic moieties required to achieve efficient gene transfer^35^. Nevertheless, cationic functionalization generally enhances the adverse cellular effects of GO alone and can compromise its colloidal stability. We have shown here that exposure to GO:siRNA complexes without cationic functionalization, even in the absence of FBS and at concentrations up to 50 µg/ml, did not induce cytotoxicity **(Figure S3)**. On the contrary, GO-PEI formulations have shown to compromise cell viability from concentrations as low as 5 and 10 µg/ml^10, 36^ and direct comparison of GO alone and GO-BPEI constructs, formed with branched PEI (BPEI), has also evidenced the detrimental effects of cationic functionalization for the safety of the vector^37^. BPEI also showed to induce the generation of large aggregates when mixed with otherwise easily dispersible GO in a variety of fluids including water, PBS and cell culture medium^37^.

A simplified vector based on the use of GO, without further functionalization, to directly interact with and transport the nucleic acid, as we investigated here, would bypass the limitations above. However, our results showed that, albeit the levels of siRNA internalized were comparable to those achieved by a lipid-based benchmark transfection reagent, that provided efficient gene silencing (Figure 3d), the biological activity of GO:siRNA complexes was impaired by a series of pitfalls in the intracellular performance of the vector that warrants further optimization.

GO was seen to localize in large vesicles, morphologically very different from those observed in cells exposed to lipo:siRNA complexes. These vesicles sequestered the material for more than 24 h in the apical region of the cell, with no apparent trafficking taking place towards endosome-lysosomal or extracellular compartments **(Figure S8, Video S3)**. The same pattern of uptake was found in other non-phagocytic cell lines when GO flakes of similar lateral dimensions were also dispersed in serum-free cell culture medium for the first 4 h^19^. In other studies, GO has been reported to enter cells mainly via endocytosis, although few-layer GO flakes of lateral dimensions under 10 µm can potentially pierce the cell membrane directly^38^. However, drawing general statements on the mechanisms of GO internalization is complicated due to the strong effect that cell type and the physicochemical characteristics of the material pose in such process, and the fact that the latter are not always thoroughly reported^7^. Nevertheless, it is interesting to note that despite the low endocytic capacity of the cell type used in our study (primary fibroblasts), the amount of internalized GO was sufficient to deliver siRNA levels comparable to those achieved with a benchmark transfection reagent. In addition, a number of studies have shown that altering the surface chemistry and size of GO materials can modulate the mechanism of cellular uptake^39-40^. This could be an opportunity to prevent GO uptake through such vesicles where the material is trapped and siRNA might be degraded.

Strategies to force GO escape from these intracellular structures may also be pursued if the entry mechanism cannot be altered. Endosomal entrapment is in fact a common obstacle faced by many nanoparticles and delivery systems, which contributes to poor transfection efficiencies, and a number of approaches have been designed to trigger escape and ensure cytoplasmic availability. Those include the generation of pores in the endosomal membrane, membrane fusion mechanisms and the proton-sponge effect, that attracts ions to burst the endosomal compartment based on elevated osmotic pressure^41^. In GO-based systems, a recurrent strategy exploits the photothermal properties of reduced GO to trigger endosomal burst via near-infrared irradiation^42-43^.

siRNA release form the GO lattice seems to be another pitfall in the performance of GO:siRNA vectors, based on our GO:siGLO studies (Figure 4c and Figure S7b), and may contribute to siRNA degradation inside the vesicles. Indeed, MD simulations performed in our study revealed that siRNA is adsorbed more tightly onto the GO surface in the presence of salts (Figure 2 and **Video S1** and **S2)**, likely due to protonation of the abundant carboxylic groups on GO alleviating electrostatic repulsion between vector and cargo. This may imply that upon addition of GO:siRNA complexes to the cell culture medium, the oligonucleotide becomes more tightly bound to GO, hampering its release. Ultimately, the specific environment inside the vesicles could be exploited to overcome this barrier. The characteristics of specific cellular environments, for example, reducing conditions in cancer cells, have been previously exploited to design cleavage mechanisms that ensure cargo release from delivery systems, including those based on GO^44-45^. Therefore, further study of the nature of the vesicles observed in this study is warranted, and will be crucial to fulfil the optimization of efficient GO:siRNA vectors, either avoiding vesicle entrapment or exploiting it to enhance intracellular trafficking of the cargo.

## Conclusion

We demonstrate here the potential of GO to perform as a non-viral gene delivery vector without the need to introduce cationic functionalization, based on its interactions with a double-stranded oligonucleotide (siRNA) that were demonstrated both experimentally and theoretically. GO can indeed act as a 2D platform to accommodate and transport a short, double-stranded nucleic acid and deliver it intracellularly at levels comparable to those of benchmark transfection reagents. However, the uptake mechanism of GO:siRNA complexes and the nature of the binding interaction under such conditions are likely to determine the lack of biological effect of the oligonucleotide cargo and represent a delivery challenge to overcome in the search for efficient and safe gene delivery vectors.

## Experimental

### Graphene oxide (GO)

GO was produced *in house* following the modified Hummer’s method as previously described^13-14^. Full characterization of the material has been provided in a previous study^15^ where it was termed small GO (s-GO) in order to differentiate it from larger counterparts that have not been used in the present study, and is summarized in **Table S1**. In brief, average lateral size of the material used here was 1 µm and thickness corresponded to 1 to 2 layers of GO (1-2 nm). Presence of oxygenated functionalities in form of hydroxyls, carboxyls and epoxides was confirmed via Fourier transform infrared spectroscopy (FT-IR) and quantified by thermogravimetric analysis (TGA). As expected, surface charge was strongly negative (ζ=-56.68 ±2.4 mV).

### siRNA

Complexation studies were performed with a scrambled siRNA sequence (siNEG, SI03650318, Qiagen) that has no homology for any known mammalian gene and was therefore also used as negative control in knockdown experiments as well as for some GO:siRNA internalization studies. siGLO green transfector indicator (D-001630-01, Dharmacon), a scrambled fluorescently-tagged siRNA modified with a nuclear targeting sequence, was used to assess release of the nucleic acids from GO. A specific siRNA duplex targeting the *Bcl-2* mouse gene (siBcl-2, MmBcl2 11, SI05462072, Qiagen) was used in gene knock-down studies

### Generation of GO:siRNA complexes

GO:siRNA complexes were formed by simple pipetting of siRNA with different amounts of GO to achieve the following final GO:siRNA mass ratios: 10:1, 20:1 and 50:1. Complexes were left to form for 30 min at room temperature prior to their characterization or use for siRNA transfection studies.

### Electrophoretic mobility shift assay

GO:siRNA complexes at 10:1, 20:1 and 50:1 mass ratios, containing 1 ug siRNA, were run on a 2% agarose gel in 0.5x TBE buffer, at 100V for 40 min. Free siRNA and lipofectamine:siRNA were used as negative and positive complexation controls, respectively. An amount of GO corresponding to the highest mass ratio tested was also run in the gel to exclude signal coming from the material. 1:10,000 Gelred™ (41003, Biotium) was added to the molten gel to visualize siRNA bands. Complexation studies were repeated 3 times for each condition.

### Quenching of fluorescence to assess complexation

The capacity of GO to quench the signal of fluorophores that interact with its surface was used to assess complexation. When siRNA was not already linked to a fluorophore, GO:siRNA complexes at 10:1, 20:1 and 50 :1 mass ratios containing 0.3 µg siRNA were incubated with the nucleic acid dye Gelred™ (1µl at 1,000X) for 30 min. Complexes where then diluted to a 1 ml total volume in TBE buffer and the emission spectrum was recorded on a spectrofluorometer (Cary Eclipse, Varian) at the optimal excitation wavelength (250 nm). When GO complexed siGLO, which is already conjugated to 6-carboxyfluorescein (6-FAM), the emission spectra at ex490 nm was directly recorded after the 30 min complexation at RT.

### Transmission electron microscopy (TEM)

GO:siRNA complexes in different mass ratios were formed as described before. Samples were deposited onto copper grids (Carbon film, 400 mesh copper) for 1 30 seconds, then washed with water and stained using a solution of 1% uranyl acetate. Images were acquired on a Tecnai 12 Biotwin TEM at 100 kV accelerating voltage. Images were taken with Gatan Orius SC1000 CCD camera.

### Enzymatic degradation assay

1 µg siRNA was exposed to increasing concentrations of DNAseI (Roche, UK), ranging from 0 to 2000 ng/µl) for 20 min at RT and then run on an agarose gel, to determine an enzyme concentration able to induce nucleic acid degradation. Free siRNA and GO:siRNA complexes at 10:1, 20:1 and 50:1 mass ratios, containing 1 µg siRNA, were subjected to such DNAseI concentration (20 ng/µl) followed by agarose gel electrophoresis (2% agarose, 0.5X TBE buffer, 100V, 40 min). Images of the gels were taken with G:Box (Syngene), 80 ms exposure.

### Attenuated Total Reflection Fourier-transform infrared spectroscopy (ATR-FTIR)

siRNA, GO and GO:siRNA complex at 50:1 mass ratio were freeze-dried to perform FTIR in solid state. Solid samples were analysed using a Bruker Vector apparatus, using the OPUS^NT^ software and collected 150 scans with a resolution of 4 cm^-1^.

### Molecular dynamics simulation

The Bcl-2 siRNA structure was built using the nucleic acid builder (NAB) functionality provided by AMBER 17^46^, with the RNA.OL3 forcefield^47-48^. The GO sheet (15 × 15 nm^2^) was created based on the Lerf-Klinowski model for graphene oxide surfaces^49^. Hydroxy, epoxy, carboxy, and carbonyl groups were added using in-house developed scripts to the initial graphene plane in such percentages as to reproduce properties and C/O ratio retrieved by XPS measurements of s-GO^15^ and taking into account existing correlations between the oxidation loci^50- 51^. Partial charges were taken from Kim et al.^52-53^, and derived by the AM1-BCC method where missing^54^; atom types were based on the general amber forcefield^55-56^. To refine the structure of GO and siRNA in solution, each system was solvated in TIP3P water^57^ and ions (Na+, Cl-) were added to obtain a neutral simulation box and to attain a 150, and 200 mM concentration of salts. The boxes were energy minimized, followed by a heating phase to take the temperature to 300 K in NVT conditions; next, the structures were relaxed in NPT conditions for 100 ns. Time step was 2 fs, periodic boundary conditions, Langevin thermostat and Berendsen barostat were applied and electrostatic interactions were treated with the particle mesh Ewald (PME) procedure^58^. From each equilibrated system, a siRNA and GO structure were extracted. The corresponding oligonucleotide was then placed onto the center of the GO sheet with the axis parallel to the basal plane and at a distance of 6 nm. The resulting GO:siRNA complexes were solvated and counterions were added to ensure system neutrality and desired ionic strength (150 and 200 mM NaCl). The structures obtained underwent the same computational procedure as the two components alone, with a production phase of 500 ns. In all simulations, the corners of the GO sheet were kept in place by Cartesian restraint applied to selected atoms to emulate the physical constraint of a larger GO sheet. Steered MD calculations pulling the siRNA center of mass away from the GO sheet were carried out at a constant speed of 10 m/s and with a spring constant of 10 kcal/(mol Å^2^) starting from stable siRNA:GO complexes at each ionic strength. The AMBER 17 suite of software running on GPU was employed to perform all MD simulations.

### Primary cell extraction, cell lines and culture

Mouse embryonic fibroblasts (MEFs) were extracted from E12.5 CD1 embryos following a standard protocol as described elsewhere^59^ and maintained in Dubelco’s Modified Eagles Medium (DMEM, D6429, Sigma) supplemented with 15% fetal bovine serum (FBS, 10500, Gibco, Lot 08G3057K) and 1% antibiotics (PenStrep, P4333, Sigma), in a 5% CO_2_ atmosphere at 37°C. They were used for a maximum of three passages. 4T1 cells were obtained from ATCC (CRL2539™) and cultured in Roswell Park Memorial Institute (RPMI) 1640 (R8758, Sigma) supplemented with 10% FBS and 1% antibiotics.

### GO:siRNA transfection experiments

Cells were exposed to GO:siRNA complexes when 70% confluency was reached. In brief, cell culture medium was removed and GO:siRNA complexes were added in fresh medium in the presence or absence of 15% FBS. When FBS-free conditions were used, complete medium supplemented with 30% FBS was added 4 hours after the initial exposure to achieve a final concentration of 15% FBS. Cytotoxicity, internalization, siRNA release and target knock-down where interrogated at different time points after transfection, as indicated in each specific experiment.

### Cytotoxicity studies

MEFs were exposed to GO:siRNA complexes, where siRNA was a scrambled sequence, at 10:1, 20:1 and 50:1 mass ratios and [siRNA]= 80 nM, in the absence of FBS. GO concentration corresponded to 10 μg/ml, 20 μg/ml and 50 μg/ml, respectively. Optical microscopy images (20X) were taken with a Zeiss Primo Vert microscope 24 h after transfection, just before the cells were detached from the cell culture vessels and stained with AnnexinV/Propidium iodide. Annexin V staining was performed according to manufacturer’s recommendations (Molecular Probes, Thermo Fisher Scientific). In brief, cells centrifuged at 1500 rpm for 5 min after trypsinization, then re-suspended in 100 μL Annexin binding buffer and stained with 2 μL Annexin V-Alexa Fluor^®^488 conjugate for 20 min at 15–25°C. Propidium Iodide (PI, 1 mg/mL, Sigma-Aldrich) was added shortly before the analysis to a final concentration of 1.5 μg/mL. 10, 000 cells were analysed on a BD FACSVerse™ flow cytometer using 488 nm excitation and 515 nm and 615 nm band pass filters for Annexin V and PI detection, respectively. Electronic compensation of the instrument was performed to exclude overlapping of the two emission spectra. GO alone was run in order to set up the gates including cell population for the analysis. Percentage of unstained (i.e. alive) cells was calculated.

### Confocal microscopy on live cells

MEFs were seeded in Cellview cell culture dishes (627870, Greiner Bio-One Ltd) and exposed to GO:siRNA complexes, where siRNA was a scrambled sequence or the transfector indicator siGLO, at 10:1, 20:1 and 50:1 mass ratios and [siRNA]= 80 nM, in the presence or absence of 15% FBS. 4 or 24 h after transfection, cells were stained with CellMask™ Green Plasma Membrane Stain (1:2500 in culture medium, C37608, Thermo Scientific). Nuclei were stained using Hoechst dye (2 µg/ml) for 2 h prior to the imaging. Cells were examined under a Zeiss 880 multiphoton confocal microscope using a 40X objective on confocal mode. Excitation wavelengths for the CellMask™, Hoechst and GO were 488 nm, 405nm and 591 nm, respectively. Emission maximum for the CellMask™ Green Plasma Membrane Stain was 520 nm, while emission wavelength for the GOs was 620 – 690 nm. Images were processed using Zeiss microscope software ZEN.

### Time lapse live confocal microscopy

MEFs were seeded in Cellview cell culture dish (627870, Greiner Bio-One Ltd) and treated with GO:siRNA at a 50:1 mass ratio, [siRNA]=80 nM, in the absence of FBS for the first 4 h. After 24 h of treatment cells were stained using CellMask™ Green Plasma Membrane Stain as described above. Time lapse live cell imaging was set for the duration of one hour. Cells were examined under a Zeiss 780 multiphoton confocal microscope using 40 x objective with a time lapse mode. Excitation wavelengths for the CellMask™ Green Plasma Membrane Stain and GOs were 488 nm and 591 nm, respectively. Emission maximum for the CellMask™ Green Plasma Membrane Stain was 520 nm, while emission wavelength for the GOs was 620 – 690 nm. Time lapse video was processed using Zeiss microscope software ZEN.

### Intracellular siRNA extraction and quantification

MEFs were transfected with GO:siBcl-2 complexes at different mass ratios, in the presence or absence of FBS. When transfection was performed in the absence of FBS, complete medium was added 4 h after initial exposure. Cells were lysed at different time points after transfection, namely 4, 12 and 24 h, and RNA was extracted with AllPrep DNA/RNA/miRNA Universal Kit (Qiagen, UK). siBcl-2 amplification was performed by stem-loop PCR using a sequence-specific TaqMan^®^ Small RNA Assays kit (Thermofisher, UK) and following manufacturer’s guidelines. In brief, RNA denaturation was performed incubating 10ng RNA with stem loop primers provided in the kit (5 min at 85^°^C, 5 min at 60^°^C). Reverse transcription was performed using TaqMan^®^ MicroRNA Reverse Transcription Kit (Thermofisher, UK) according to the following protocol: 16^°^C for 30 minutes, 42^°^C for 30 minutes and 85^°^C for 5 minutes. qPCR amplification was performed with TaqMan^®^ Small RNA Assays and TaqMan^®^ Universal PCR Master Mix II no UNG (Thermofisher, UK), as follows: 50^°^C for 2 minutes, 95^°^C for 10 minutes and 40 cycles of 95^°^C for 15 seconds and 60^°^C for 1 minute. A calibration curve was built with siBcl-2 to calculate the amount of intracellular siBcl-2.

### RNA extraction and real-time RT-PCR

24 h after transfection with GO:siRNA complexes, cells were lysed and total RNA was extracted with Purelink^®^ RNA Mini Kit (12183025, Applied Biosystems) following the manufacturer’s guidelines. RNA quantity and quality were assessed by spectroscopy with a Biophotometer (Eppendorf). All samples included in the study had A260/A280 and A260/A230 ratios greater than 1.8. 1 ug RNA was used for each cDNA synthesis reaction, which was performed with High Capacity Reverse Transcription kit (4368814, Applied Biosystems) according to the following protocol: 10 min at 25^°^C, 120 min at 37^°^C and 5 min at 85^°^C, followed by cooling at 4^°^C. 2 ul of the resulting cDNA were used in each qPCR reaction, which was performed with SYBR green chemistry (PowerUp™ SYBR™ Green Master Mix, A25742, Applied Biosystems) according to the following protocol: 2 min at 50^°^C, 2 min at 95^°^C, (15 sec at 95^°^C and 1 min at 60^°^C) × 40 cycles; and followed by a melt curve analysis as specified by the manufacturer to confirm amplification of a single product. Primer sequences for the amplification of Bcl-2 mRNA were as follows: Fwd 5’GAACTGGGGGAGGATTGTGG3’, Rv 5’GCATGCTGGGGCCATATAGT3’. Primer pairs for reference genes are listed in **Supplementary Table 1**. Data was normalized by the Livak method using two reference genes, *Hmbs* and *Tbp*, validated by the geNorm algorithm integrated in qbase+ software. Gene expression levels were normalized to those of untreated cells.

### Validation of reference genes for real-time RT-qPCR normalization

Ten candidate reference genes with different cell functions **(Supplementary table 1)** were evaluated for the stability of their expression levels under the specific experimental conditions in this study. In brief, MEFs were transfected with the following conditions: untreated, Lipofectamine:siBlc-2, GO:siBcl-2 (20:1), GO:siBcl-2 (50:1) and GO alone (equivalent concentration to that in the 50:1 mass ratio). [siBcl-2] was 80 nM in all conditions. Transfection was performed in absence of FBS for the first 4 h after which FBS was added to reach a final 15% FBS concentration. mRNA was extracted 24 h after transfection and real-time RT-qPCR was performed as above. The geNorm algorithm as described elsewhere^60^ integrated in qbase+ software was used to calculate the stability factor of each gene (geNorm M, **Figure S4a**) and the optimal number of reference genes to be included in the study (geNorm V, **Figure S4b**).

## Supporting information

## Acknowledgements

This work was funded by Working Package 5 of the Graphene Flagship (European Commission). The authors would like to acknowledge the staff in the Faculty of Biology, Medicine and Health Electron Microscopy Facility for their assistance and the Wellcome Trust for equipment grant support to such facility. The authors would also like to thank Dr David Spiller for his assistance with confocal fluorescence life imaging. DM and PP acknowledge support by the SIR Program of the Italian Ministry of University Research under award number RBSI14PBC6.

## Author contributions

IdL and KK initiated and designed the study. IdL lead the project and performed the majority of experiments. SV performed live-cell imaging. MB performed siBcl-2 quantification. DM and PP performed MD simulations. MM obtained TEM images. AEA contributed to FTIR data and AFR produced and characterized the GO material used in the study. IdL, SV, PP and KK wrote the manuscript.

