## Supplementary material for "Graphene Oxide as 2D Platform for Complexation and Intracellular Delivery of siRNA"

### Supporting Figure 1

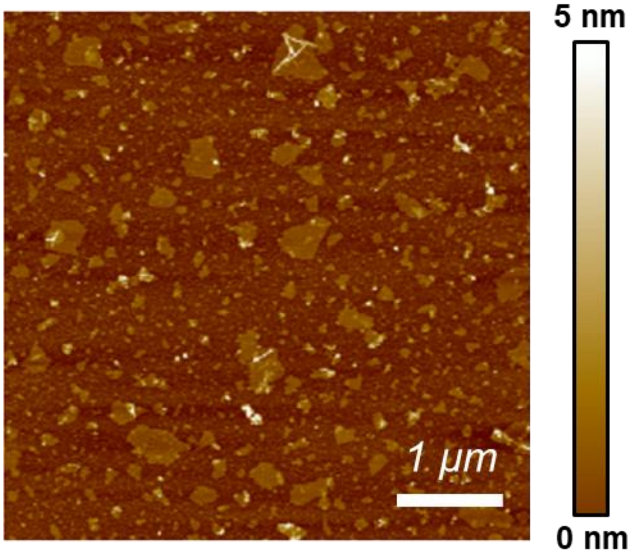


**Figure S1. AFM image of GO flakes.** AFM analysis indicated a distribution of GO flakes with lateral dimensions in the 0.05 – 0.5 μm range and thickness within 1.4 ± 0.5 nm, equivalent to 1-2 layers of GO.

### Supporting Figure 2

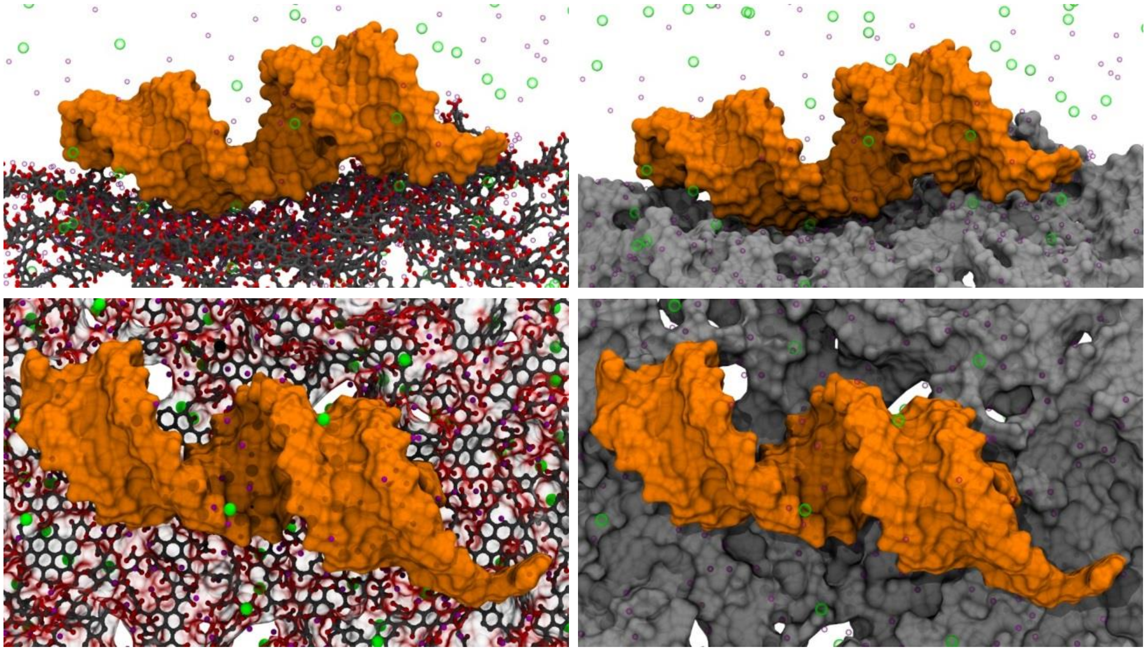

**Figure S2. GO:siRNA binding by MD simulation.** Snapshots extracted from MD calculation of siRNA (represented as an orange shaded surface) in complex with a GO sheet at 200 mM NaCl ionic strength (top panels, lateral view; bottom panels, top view). In the left panels, the chemical structure of GO is explicitly represented with carbon and oxygen atoms colored in grey and red, respectively. In the right panels, we depicted the GO sheet as a grey surface to highlight 3D features. To preserve clarity, water is not shown and $Na^+$ and $Cl^-$ ions are visualized as purple and green spheres, respectively.

### Supporting Figure 3

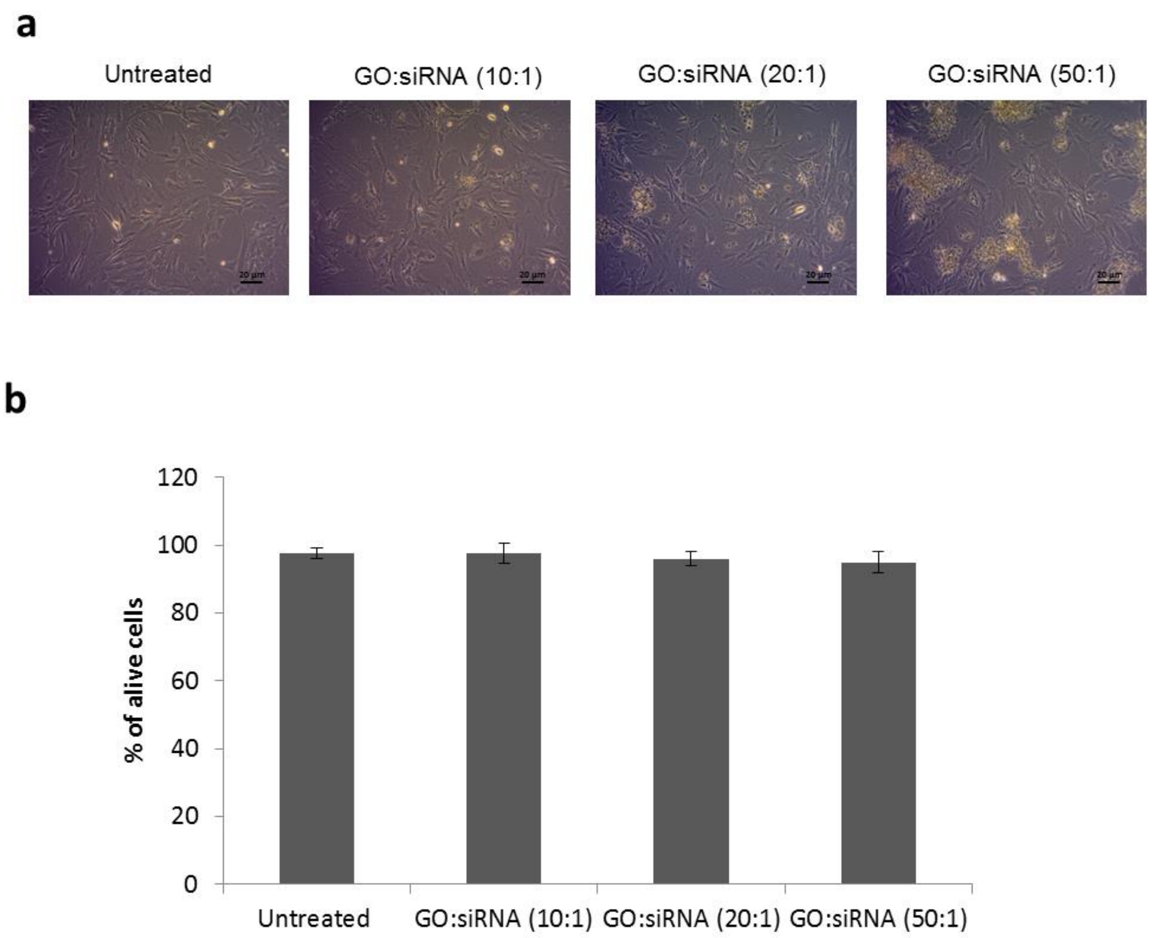


**Figure S3. Effects of GO:siRNA transfection on cell viability. (a)** Optical microscopy images of MEF exposed to GO:siRNA complexes for 24 h. **(b)** Cell viability assessed by flow cytometry (AnnexinV/PI staining).

### Supporting Figure 4

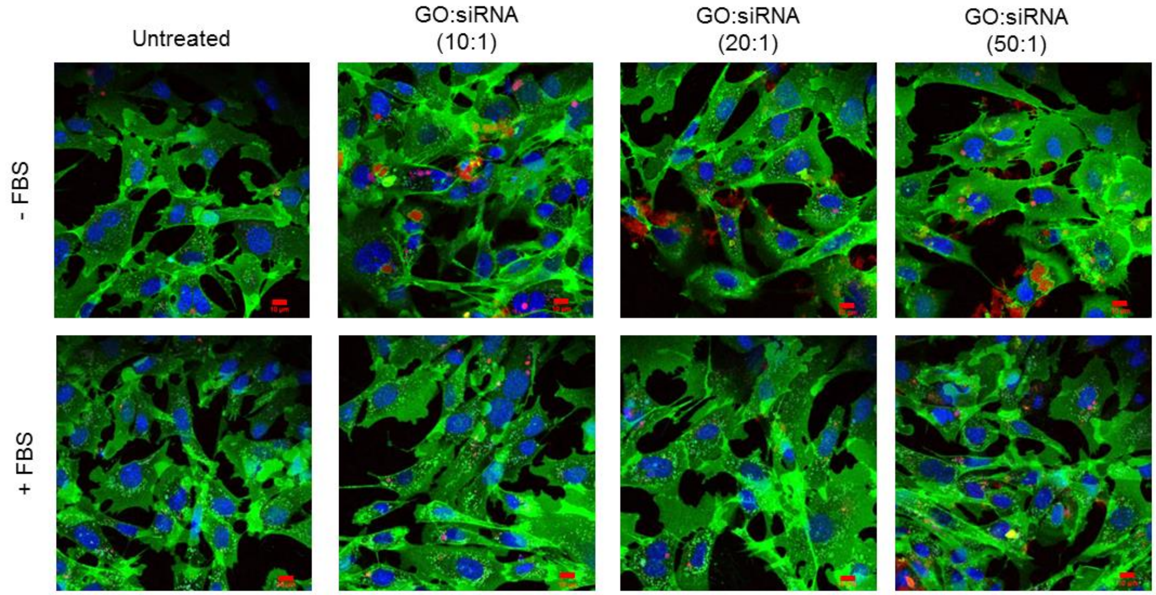


**Figure S4. Uptake of GO:siRNA complexes in the presence of absence of serum proteins.** GO uptake in MEFs was visualized by confocal microscopy, thanks to the intrinsic fluorescence of the material, 24 h after exposure. The upper panel represents serum-free treatment conditions, while the lower panel shows that the presence of 15% FBS in the medium significantly impaired GO uptake. Green – Cell Membrane Mask, Red – GO, Blue – nucleus (Hoechst). Scale bar shows 10 µm.

### Supporting Figure 5

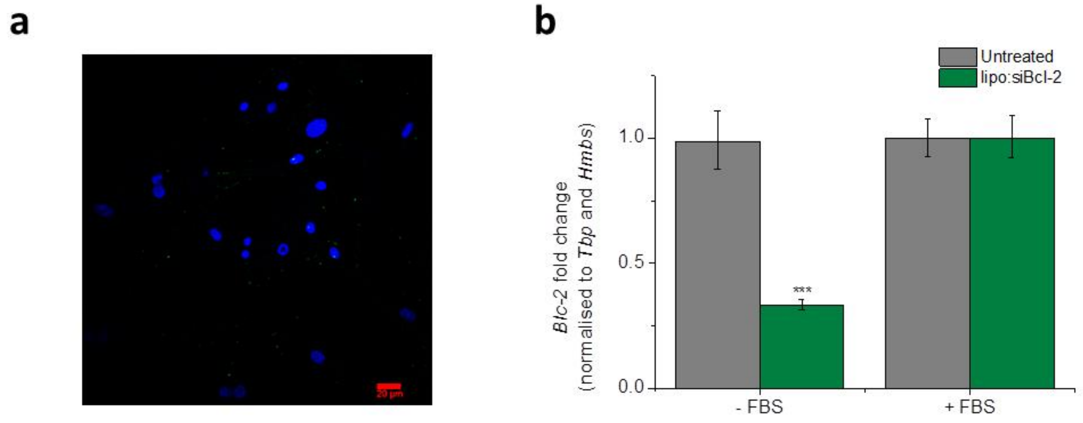


**Figure S5. FBS blocks siRNA delivery with lipo:siRNA complexes. (a)** Confocal microscopy of MEFs, 4 h after transfection with lipo:siGLO in the presence of FBS, indicated absence of siRNA intracellular delivery. Scale bar shows 20 µm. **(b)** Analysis of *Bcl-2* mRNA levels by RT-qPCR 24 h after transfection with lipo:siBcl-2 in the presence or absence of FBS confirms the absence of target downregulation in the presence of serum proteins (***p<0.001, one-way ANOVA and Tukey post-hoc test, n=3).

### Supporting Figure 6

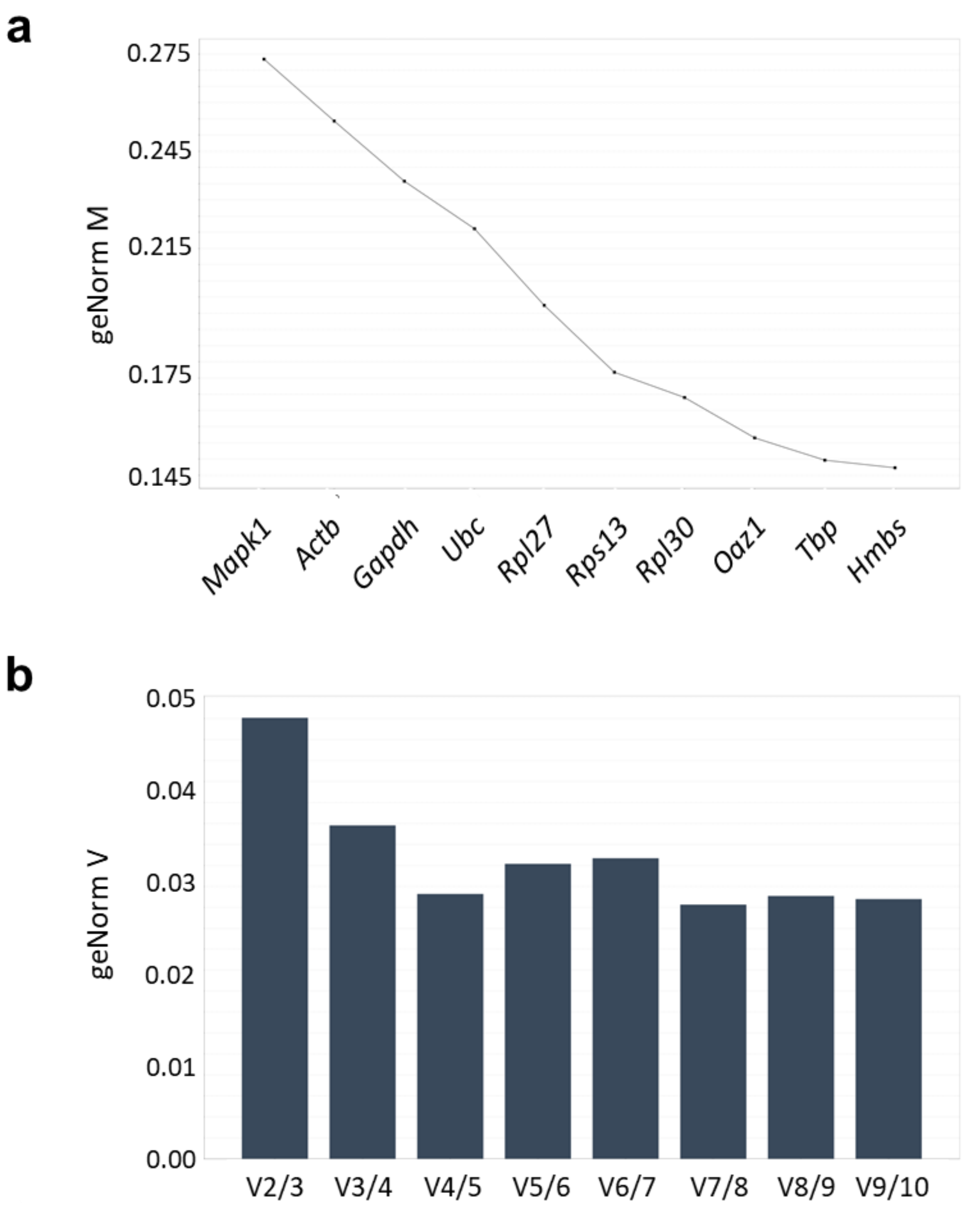


**Figure S6. Analysis of reference gene expression stability upon transfection with GO:siRNA vectors.** Expression of ten candidate reference genes was analyzed by RT-qPCR and GeNorm software. **(a)** Stability values for each candidate reference gene (GeNorm M). **(b)** Pair-wise comparison of all reference genes indicated that two genes should be included as reference for reliable normalization of gene expression data (GeNorm V).

### Supporting Figure 7

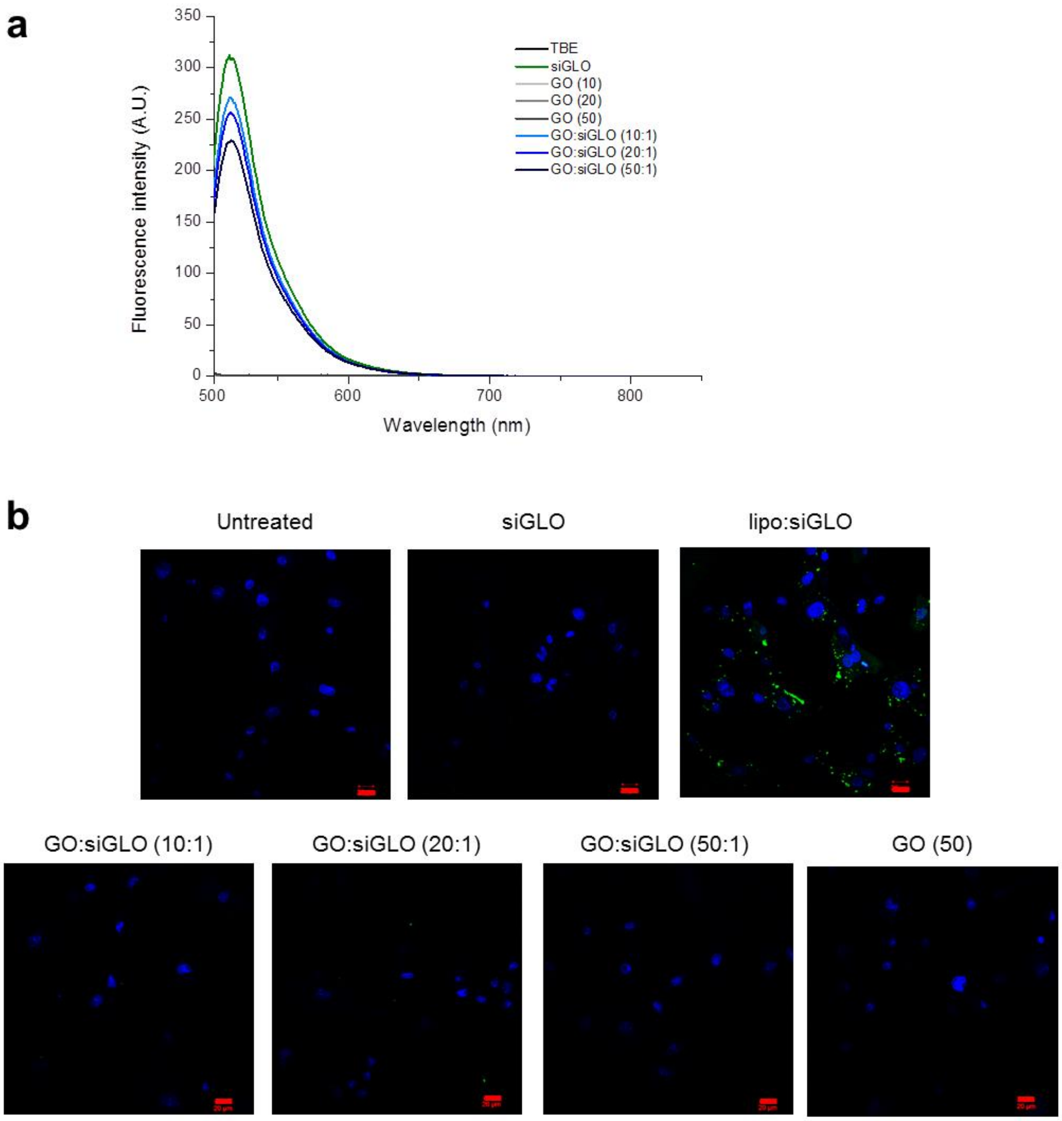


**Figure S7. Complexation and intracellular delivery of a nuclear-targeted, fluorescently labelled siRNA (siGLO). (a)** GO:siGLO complexation was confirmed by a decrease in fluorescence signal with increasing GO:siRNA mass rations. **(b)** 24 h after transfection, most of the signal had disappeared from the lipo:siGLO condition. No signal was detected in GO:siGLO condition. Green – 6-FAM labelling, siGLO oligonucleotide, Blue – nucleus (Hoechst). Scale bar shows 20 µm.

### Supporting Figure 8

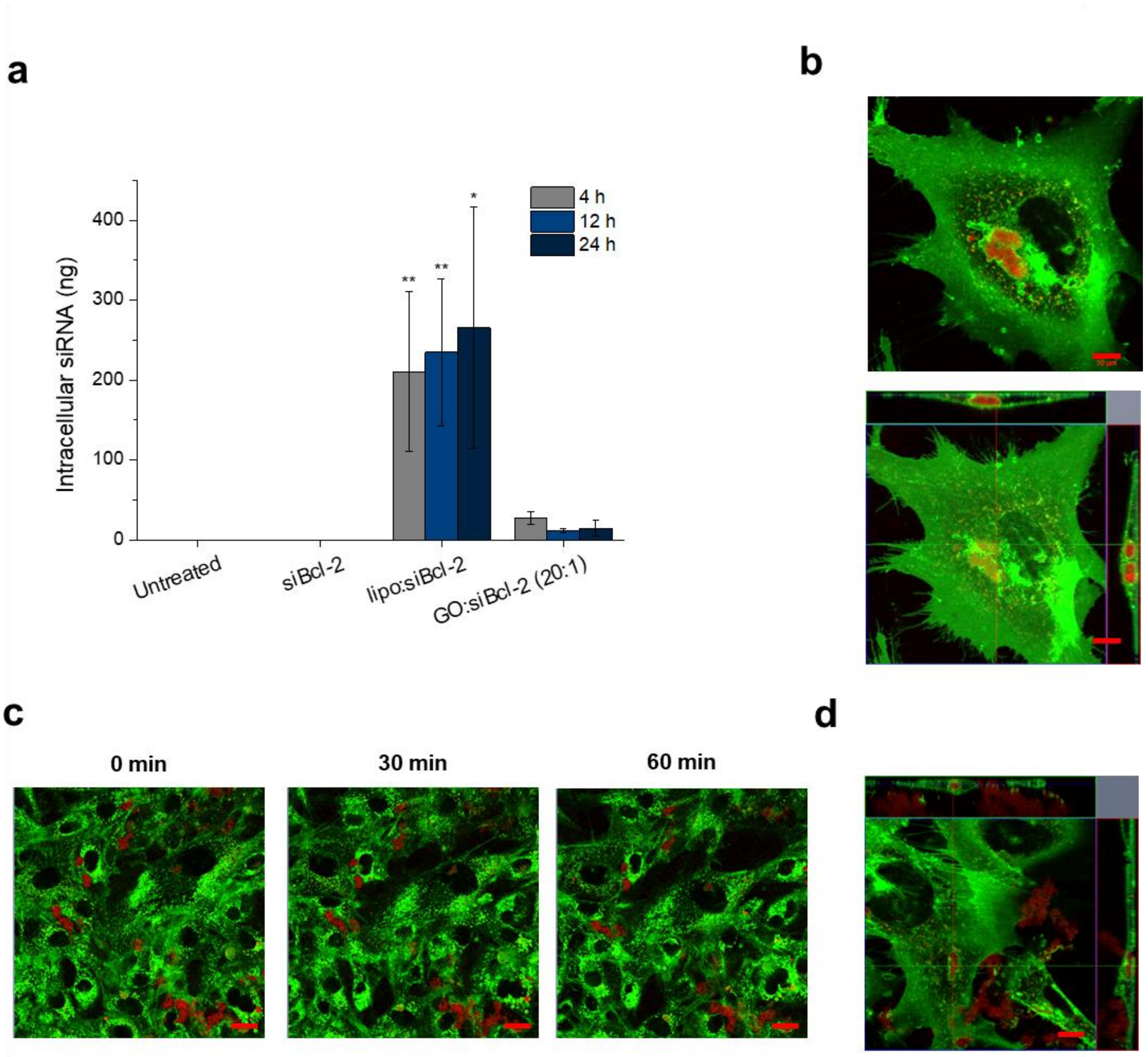


**Figure S8. Quantification of intracellularly delivered siRNA in 4T1 cells. (a)** A PCR-based method was used to determine the amount of siRNA successfully internalised in 4T1 cells 4, 12 and 24 h after transfection with GO:siBcl-2 complexes. (*$p<0.05$ and **$p<0.001$, one-way ANOVA and Tukey post-hoc test, n=3). **(b)** Vesicular entrapment of GO:siRNA complex already after 6 h of treatment. Left panel represents a section in the middle of the cell, while right panel shows orthogonal projection of the vesicular region. Green – Cell Membrane Mask, Red – GO. Scale bar shows 10 µm. **(c)** Captures of the video taken within an hour difference (0 , 30 and 60 min) after 24 h of treatment (using GO:siRNA at 50:1 mass ratio) show that the vesicle moves with the cell but is not further trafficked. Green – Cell Membrane Mask, Red – GO. Scale bar shows 20 µm. (d) Cells treated with uncoated GO (i.e. in the absence of siRNA), in a concentration equivalent to that of the 50:1 mass ratio, for 4 h. Green – Cell Membrane Mask, Red – GO. Scale bar shows 10 µm.

### Supporting Table 1

| | *Technique* | *Results* |
|---|---|---|
| ***Lateral dimensions**** | *AFM* | 0.050 – 0.5 μm |
| | *TEM* | 0.1 – 2 μm |
| ***Thickness**** | *AFM* | 1.4 ± 0.5 nm (1-2 layers) |
| ***Optical Properties*** | *Absorbance ($\lambda$ = 230 nm)* | $A = 0.050 * C_{GO}$ (μg/mL) |
| | *Fluorescence ($\lambda_{exc}$ = 525 nm)* | $F_{600} = 0.823 * C_{GO}$ (μg/mL) |
| ***Degree of defects ($I_D/I_G$)*** | *Raman spectroscopy* | 1.36 ± 0.03 |
| ***Surface charge*** | *ζ-potential* | -55.9 ± 1.4 mV |
| ***Functionalisation degree*** | *TGA* | 41% |
| ***Chemical composition (Purity)*** | *XPS* | C: 67.6%, O: 32.2%, (99.8%) S: 0.2% |
| ***C:O ratio*** | *XPS* | 2.1 |
| ***π-π, O=C-O, C=O, C-O-C, C-C & C=C*** | *XPS* | 3.0%, 15.3%, 31.2%, 11.3%, 39.2% |

**Table S1. Physicochemical characteristics of GO used in this study.** *Lateral dimensions and thickness are reported as a range between the minimum and maximum sizes detected. Full characterization of the material is provided in Mukherjee et al[15]. This Table has been adapted from the original publication.

### Supporting Table 2

**Table S2.** Primer sequences used in the reference gene stability study.

| *Gene symbol* | *Gene name* | *Gene ID* | *Accession number (mRNA)* | *Cell function* | *Fwd Seq* | *Tm* | *Rv seq* | *Tm* | *Amplicon size* |
|---|---|---|---|---|---|---|---|---|---|
| ***Rps13*** | Ribosomal protein S13 | 68052 | NM_026533.3 | Translation | CCCAGGTCCGTTTTGTGACT | 60 | TCCTCTCAAGGTGCTTTCGG | 60 | 132 |
| ***Rpl27*** | Ribosomal protein L27 | 19942 | NM_011289.3 | Translation | CAAAAACGCAGTGCCCGA | 59 | CTTACGGAGAGGTGGCTTCA | 59 | 120 |
| ***Rpl30*** | Ribosomal protein L30 | 19946 | NM_009083.4<br>NM_001163485.1 | Translation | GAAGAGCTTTGCATTGTGGGAG | 60 | CCATCTTCCTGCCTTAGGTGC | 61 | 102 |
| ***OAZ1*** | Ornithine decarboxylase antizyme 1 | 18245 | NM_008753.4<br>NM_001301034.1 | Metabolism | GGGTTGCCCTTAATTGCTGT | 59 | TCTTGTCGTTAGACGTCGGC | 60 | 187 |
| ***Actb*** | β-actin | 11461 | NM_007393.5 | Structural | CTGAGCTGCGTTTTACACCC | 59 | CGCCTTCACCGTTCCAGTTT | 61 | 200 |
| ***Gapdh*** | glyceraldehyde-3 phosphate dehydrogenase | 14433 | NM_001289726.1<br>NM_008084.3<br>XM_017321385.1 | Metabolism | AGGTCGGTGTGAACGGATTTG | 61 | TGTAGACCATGTAGTTGAGGTCA | 59 | 123 |
| ***Mapk1*** | mitogen-activated protein kinase 1 | 26413 | NM_011949.3<br>NM_001038663.1<br>XM_006522147.3 | Signalling | GGTTGTTCCCAAATGCTGACT | 59 | CAACTTCAATCCTCTTGTGAGGG | 59 | 84 |
| ***Ubc*** | Ubiquitin C | 22190 | NM_019639.4 | Metabolism | CAAACAGGAAGACAGACGTACC | 59 | CCCATCACACCCAAGAACAAG | 59 | 80 |
| ***Hmbs*** | Hydroxymethylbilane synthase | 15288 | NM_013551.2<br>NM_001110251.1 | Metabolism | ATCTTGGACCTAGTGAGTGTGT | 58 | GTACAGTTGCCCATCTTTCATCA | 59 | 141 |
| ***Tbp*** | TATA-box binding protein | 21374 | NM_013684.3 | Transcription | TTTGGCTAGGTTTCTGCGGT | 60 | GCCCTGAGCATAAGGTGGAA | 60 | 195 |

#### Snapshot from Supporting Video 1

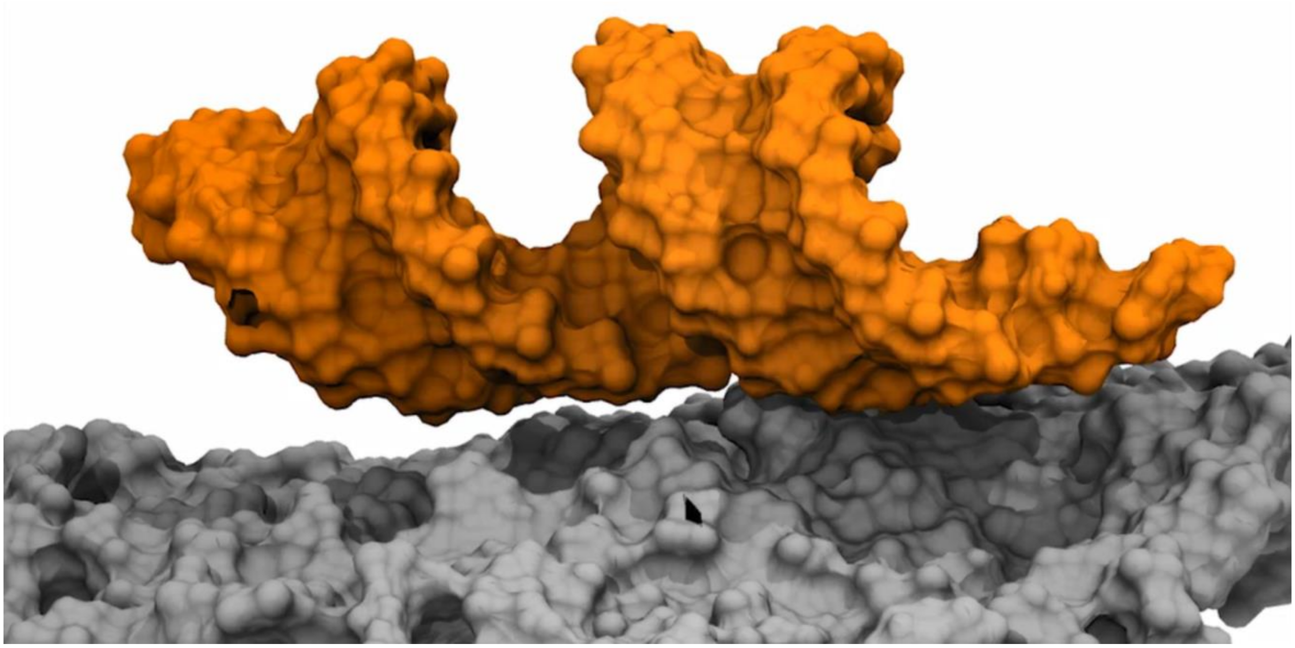

**Video S1. GO:siRNA binding by MD simulation without complementary salts.** Movie extracted from MD simulation of GO complexing siRNA in aqueous solution at 0 nM NaCl ionic strength.

#### Snapshot from Supporting Video 2

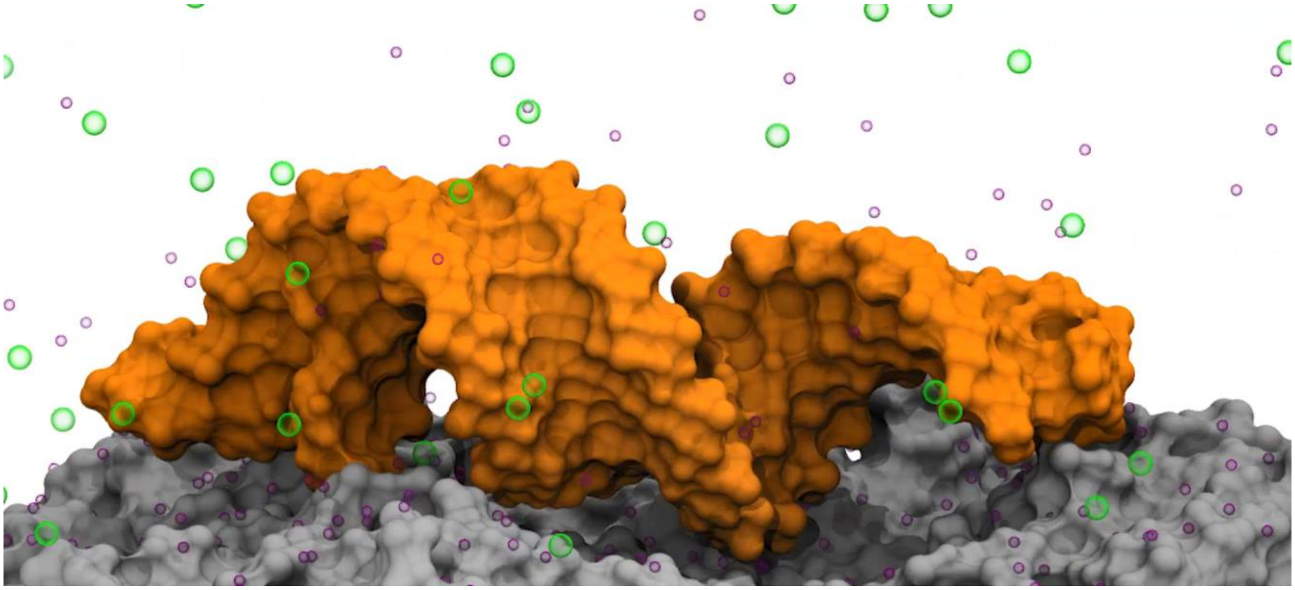

**Video S2. GO:siRNA binding by MD simulation at 150 mM NaCl.** Movie extracted from MD simulation of GO complexing siRNA in aqueous solution at 150 mM NaCl ionic strength.

#### Snapshot from Supporting Video 3

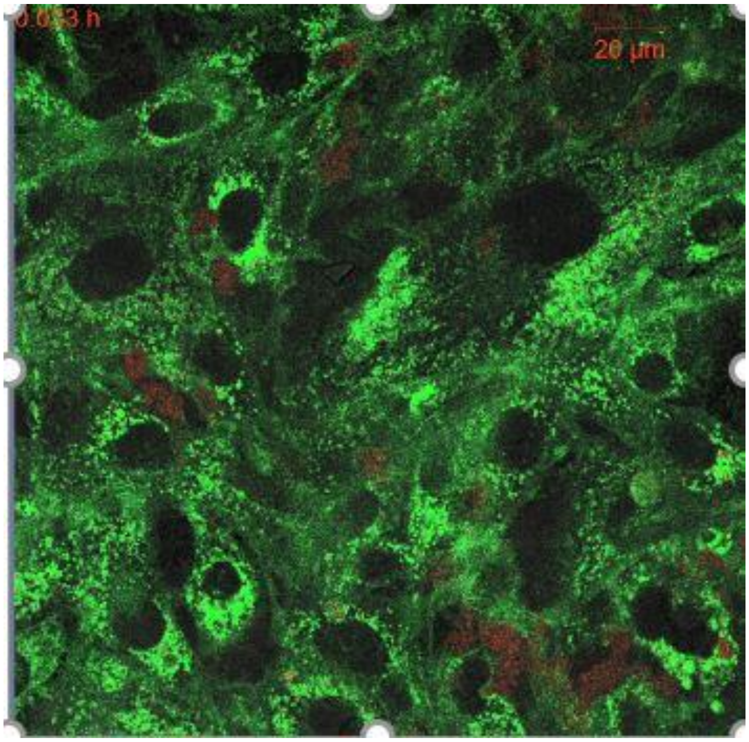


**Video S3. Intracellular fate of GO:siRNA complexes.** An intracellular vesicle containing GO was followed for 1 h, 24 h after transfection with GO:siRNA (50:1). Staining of the cells is as follows: Green – Cell Membrane Mask, Red – GO.
